# Astrocytic Regulation of Cocaine Locomotor Sensitization in EcoHIV Infected Mice

**DOI:** 10.1101/2024.09.04.611213

**Authors:** Qiaowei Xie, Rohan Dasari, Mark D. Namba, Lauren A. Buck, Christine M. Side, Kyewon Park, Joshua G. Jackson, Jacqueline M. Barker

## Abstract

Cocaine use disorder (CUD) is highly comorbid with HIV infection and worsens HIV outcomes. Preclinical research on the outcomes of HIV infection may yield crucial information on neurobehavioral changes resulting from chronic drug exposure in people living with HIV (PLWH). Repeated exposure to cocaine alters behavioral responses to cocaine. This includes development of cocaine locomotor sensitization – or increased locomotor responses to the same doses of cocaine - which depends on nucleus accumbens (NAc) neural plasticity. NAc astrocytes are key regulators of neural activity and plasticity, and their function can be impaired by cocaine exposure and HIV infection, thus implicating them as potential regulators of HIV-induced changes in behavioral response to cocaine. To characterize the effects of HIV infection on cocaine locomotor sensitization, we employed the EcoHIV mouse model to assess changes in locomotor responses after repeated cocaine (10mg/kg) exposure and challenge. EcoHIV infection potentiated expression of cocaine sensitization. We also identified EcoHIV-induced increases in expression of the astrocytic nuclear marker Sox9 selectively in the NAc core. To investigate whether modulation of NAc astrocytes could reverse EcoHIV-induced deficits, we employed a chemogenetic approach. We found that chemogenetic activation of NAc astrocyte Gq signaling attenuated EcoHIV-enhanced cocaine sensitization. We propose that HIV infection contributes to cocaine behavioral sensitization and induces adaptations in NAc astrocytes, while promoting NAc astrocytic Gq-signaling can recover EcoHIV-induced behavioral changes. These findings identify potential cellular substrates of disordered cocaine-driven behavior in the context of HIV infection and point toward strategies to reduce cocaine-related behavior in PLWH.

## Introduction

Human immunodeficiency virus (HIV) impairs the central nervous system (CNS) during chronic infection, creating risk for comorbidities and cognitive impairment. One common comorbidity is the use of psychoactive drugs, including psychostimulants such as cocaine, which is more prevalent amongst people living with HIV (PLWH) than in the general population (Mimiaga et al., 2013). Cocaine use disorder (CUD) can interact with HIV to exacerbate neuropathogenesis and cognitive impairment (Baum et al., 2009; Buch et al., 2011; Dash et al., 2015) and alter neuronal function within reward circuitry (Meade et al., 2018), necessitating effective therapeutic strategies in PLWH. Despite the high prevalence of co-occurring CUD and HIV infection, the mechanisms by which drug exposure and HIV infection interact to alter behavioral responses are still unclear.

Both cocaine and HIV are independently associated with deficits in brain reward processing regions. Cocaine exposure alters neuronal plasticity within the nucleus accumbens (NAc), which is a key substrate of cocaine-related behaviors including locomotor response, reward-learning and processing, and relapse-like behavior (Koya et al., 2009; Koob and Volkow, 2010, 2016; Perrine et al., 2015). Locomotor sensitization is a behavioral adaptation to psychostimulant drugs such as cocaine in which the cocaine-induced locomotor response becomes more robust upon repeated exposure to the same dose, i.e., is sensitized (Van Zessen et al., 2021). It has been documented that behavioral sensitization to cocaine reflects molecular and neuronal adaptions in reward circuits that contribute to cocaine-associated pathophysiological emotional state and the development of cocaine seeking behaviors (Parsons and Justice, 1993; Thomas et al., 2001; Crombag et al., 2002; Steketee and Kalivas, 2011; Allain et al., 2017). Specifically, cocaine-induced locomotor sensitization is mediated by the neuroadaptations within the NAc, including upregulation of glutamate receptor expression and priming the long term potentiation (LTP) in the NAc (Boudreau et al., 2007; Brown et al., 2011).

Astrocytes play an important role in regulating neurotransmission and plasticity, and are implicated in the regulation of drug-related behavior (Sofroniew and Vinters, 2010; Scofield et al., 2015; Bazargani and Attwell, 2016). Chronic cocaine exposure elicits morphological, molecular, and functional changes in astrocytes that contribute to synaptic plasticity and behavioral alterations (Scofield et al., 2016; Kim et al., 2018; O’Donovan et al., 2021; Wang et al., 2021b). For example, chronic cocaine exposure induces astrocyte-specific functional change via a series of calcium (Ca^2+^) responses that promote neuronal adaptations in response to cocaine seeking and relapse (O’Donovan et al., 2021). Astrocytes are also impacted by CNS HIV infection and may serve as a substrate mediating interactive HIV and cocaine effects. HIV-1 enters the CNS through infiltration of infected peripheral monocytes and macrophage where it can infect brain resident glia cells including microglia and astrocytes to establish virial reservoir (Wallet et al., 2019; Namba et al., 2023). Although HIV-1 infection in astrocytes is thought to be non-productive, HIV-1 infection and its viral proteins induce neuroinflammation responses and promote astrocyte reactivity, which participates in driving neuronal dysfunction and cognitive impairment in PLWH. Increased astrocyte reactivity is associated with potentiated glutamate transmission on the reward circuit in response to cocaine exposure (Satarker et al., 2022). Thus, HIV infection could augment the effect of cocaine on astrocyte adaptions that contribute to enhanced locomotor sensitivity to cocaine.

Here, we investigate the role of astrocytes in modulating cocaine locomotor sensitization in the context of HIV infection. Others have found that chemogenetic modulation of astrocyte Gq signaling showed an inhibitory influence on drug seeking. For example, astrocytic Gq activation in the NAc suppressed cue-induced reinstatement of cocaine seeking (Scofield et al., 2015) and ethanol self-administration (Bull et al., 2014). These studies have shown activation of NAc astrocyte via Gq DREADD improves astrocytic glutamatergic regulation that results in attenuation of relapse propensity and drug self-administration. However, the effect of NAc astrocytic Gq activation on cocaine locomotor sensitization – independently or in a model of co-occurring HIV infection - has not to date been characterized.

We hypothesized that HIV infection interacts with cocaine to disrupt astrocytes in the NAc and exacerbate cocaine locomotor sensitization in a mouse model of HIV infection. To test this, we used the EcoHIV mouse model, employing a chimeric HIV-1 virus that infects rodents, and characterized changes in cocaine locomotor sensitization and expression of astrocyte markers. We further investigated whether modulating astrocyte activity using chemogenetics altered the expression of locomotor sensitization in EcoHIV-infected mice. The results indicate that EcoHIV infection heightened cocaine locomotor sensitization and perturbed of NAc astrocytes, and further that NAc astrocytic signaling can be targeted to rescue cocaine-related behaviors in EcoHIV-infected mice, suggesting that astrocytes are a promising target for recovery of infection-induced behavioral changes.

## Materials and Methods

### Subjects

Adult male (n = 50) and female (n = 50) C57BL/6J mice (9 weeks of age upon arrival) were obtained from Jackson Laboratories. Following arrival, mice were group housed in same-sex cages for 7 days to acclimatize with *ad libitum* access to a standard chow diet and water. Mice were housed at the Drexel University College of Medicine under standard 12-hour light:12-hour dark conditions in microisolation conditions throughout the experiments. All experiments were approved by the Institutional Animal Care and Use Committee at Drexel University.

### EcoHIV-NDK inoculation

Plasmid DNA encoding the EcoHIV-NDK coding sequence (gift from Dr. David Volsky) was grown overnight in Stbl2 bacterial cells (ThermoFisher#10268019) and the plasmid DNA was purified using an endotoxin free plasmid purification kit (ZymoPure #D4200). Purified DNA was transfected into nearly confluent (80–90%)10 cm^2^ plates of low passage LentiX 293T cells (#632180, Takara Bio, San Jose, CA, USA), using a calcium phosphate transfection kit (Takara #631312). The cell culture supernatant was collected at 48 hours post-transfection using centrifugation at low speed (1500×g, 4°C), followed by passage through a cell strainer (40μm) to remove LentiX cells and cell debris. The supernatant, containing viral particles, was mixed 4:1 with a homemade lentiviral concentrator solution (4X; MD Anderson) composed of 40% (w/v) PEG-8000 and 1.2 M NaCl in PBS (pH 7.4). The supernatant–PEG mixture was incubated overnight at 4 °C on an orbital shaker (60 rpm). The mixture was centrifuged at 1500×g for 30 min at 4 °C. After centrifugation, the medium was removed, and the viral pellet was resuspended in cold, sterile PBS. The viral titer (p24 core antigen content) was determined initially using a LentiX GoStix Plus titration kit (#631280, Takara Bio, San Jose, CA, USA) and subsequently using an HIV p24 AlphaLISA detection kit (#AL291C, PerkinElmer, Waltman, MA, USA). Viral stocks were aliquoted and stored at−80 °C upon usage.

Following one week of acclimation, adult male and female mice were inoculated with 300 ng p24 equivalent EcoHIV-NDK or PBS (virus culture vehicle) as sham control by i.p. injection. To ensure housing consistency, all mice were singly housed. Blood samples were collected from all mice 1, 3, and 5 weeks following EcoHIV inoculation. Five weeks after inoculation, mice were assigned for behavioral training. This dose of virus and length of infection was selected as it produces systemic infection and immune response in the CNS and periphery (Potash et al., 2005; Kelschenbach et al., 2012, 2019; Gu et al., 2018; Xie et al., 2024). Following the completion of all experiments, to confirm the terminal EcoHIV-NDK infection status, spleens were isolated and flash-frozen on dry ice prior to perfusion and stored at -80C until processing. Splenic viral DNA burden was measured using Qiagen QIAamp DNA Mini Kit (#51304, Qiagen, Germantown, MD, US). Viral DNA was analyzed by the University of Pennsylvania Center for AIDS Research (CFAR). qPCR was conducted as described (Xie et al., 2024) using primers that amply sequences with HIV-LTR provided below. The OD value was detected by a NanoDrop™ spectrophotometer (Thermo Scientific) and used to determine the input cell numbers to normalize the data.

Kumar LTR F, GCCTCAATAAAGCTTGCCTTGA

Kumar LTR R, GGGCGCCACTGCTAGAGA

Kumar LTR Probe (FAM/BHQ), 5’CCAGAGTCACACAACAGACGGGCACA 3’

### Cocaine locomotor sensitization

Locomotor activity was assessed in transparent open field chambers (Med Associates). Each chamber contained a 16 X 16 photobeam array. Photobeam breaks were automatically recorded with Med Associates system. **Figure 1A** provides a timeline of the behavioral procedure.

**Figure 1.**
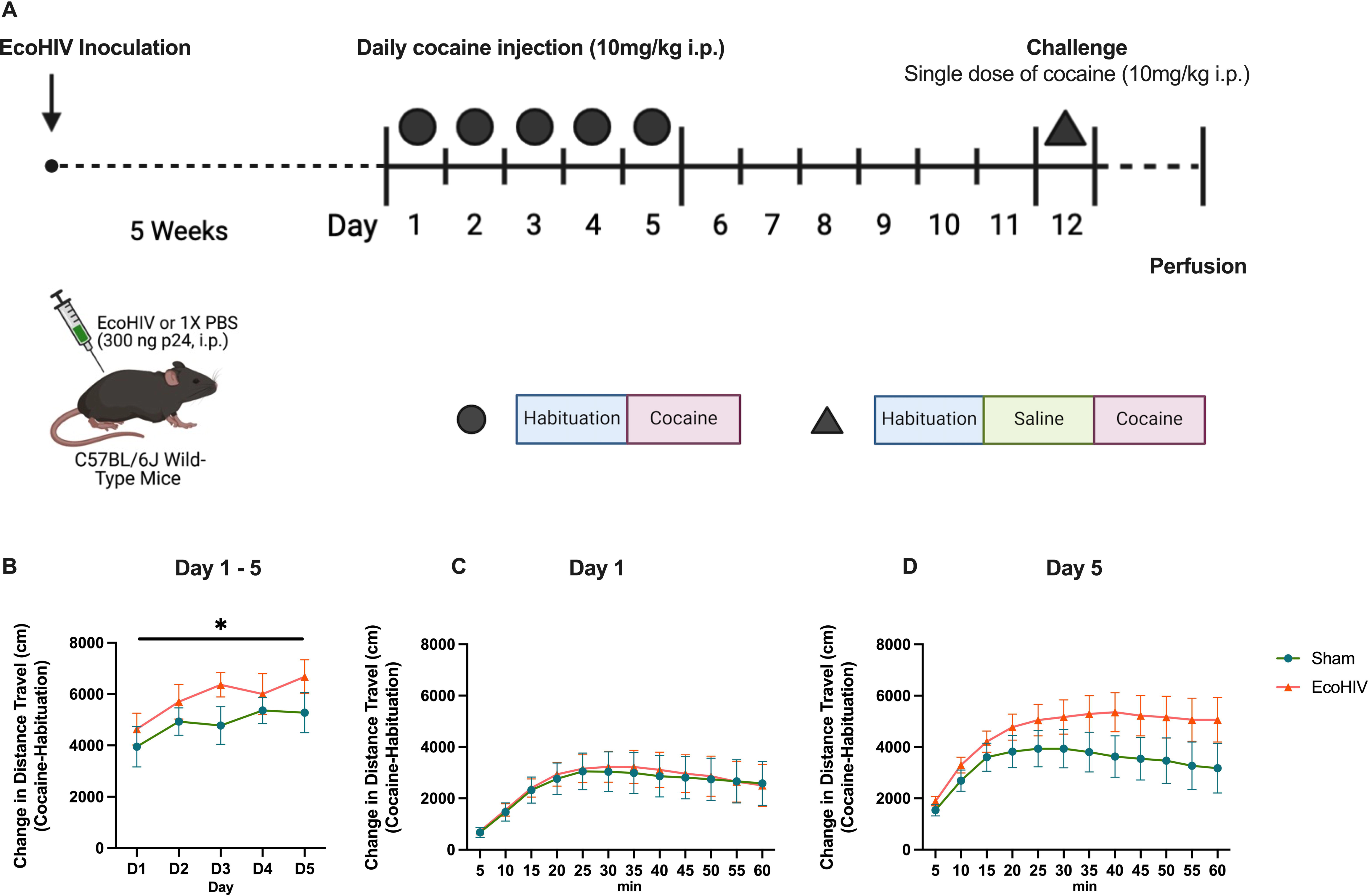
Induction of cocaine locomotor sensitization. (**A**) Timeline of experiments. Mice underwent EcoHIV or sham inoculation 5 weeks prior to undergoing a cocaine locomotor sensitization protocol. Sensitization consisted of 5 daily injections of cocaine (10 mg/kg), followed by 6 days of abstinence and a final challenge dose of cocaine. (**B**) Distance traveled was increased in both sham (n=18) and EcoHIV-infected mice (n=17) in response to the repeated cocaine injection. No effect of EcoHIV was detected across the daily injections. To determine if the time course of locomotor activity was impacted by EcoHIV infection, the change in distance traveled during the 60 min open field sessions in 5-min bins on the first (day 1) and final (day 5) session of daily cocaine exposure were analyzed. No effect of EcoHIV was observed on Day 1 (**C**). Following 5 daily cocaine injections, EcoHIV-infected mice showed a trend toward higher cocaine locomotor responses after 5 daily exposures to cocaine (**D**). Timeline created with Biorender.com. Data represent mean +/- SEM. *p < 0.05.

Mice were habituated to the test room at least 1 h before initiation of the experiment. After habituation, mice were placed individually into an open field chamber. Baseline locomotor activity was assessed in the chamber for 60 min. Mice then received an injection of cocaine hydrochloride (10 mg/kg, i.p.) and were returned to the locomotor chamber. Cocaine-induced locomotor activity was recorded for 60 min. Measures included distance traveled (cm), ambulatory counts (defined as: photobeam breaks during travel), ambulatory episodes (defined as: individual movement bouts) average velocity, and stereotypy counts (defined as: photobeam breaks without ambulation/travel). Locomotor measurements were registered in 5-min bins. Total distance traveled during the first 30 min of habituation and first 30 min after cocaine injection were analyzed in the EcoHIV-infected and sham control groups.

The cocaine injections were repeated daily at the same time of day for 5 consecutive days. Mice then underwent 6 days of abstinence followed by a 3-hour challenge test. In the challenge test, mice were habituated for 60 min, followed by 60 min monitoring after a saline injection to exclude injection-specific responses. In the cohort of mice used for chemogenetic studies, a subset of mice received clozapine-N-oxide in lieu of saline (details below). Mice then received a challenge dose of cocaine (10 mg/kg, i.p.) and were returned to the chamber for an additional 60 min. The change in distance traveled was calculated by subtracting the distance traveled in the first 30 minutes after the saline injection from the distance traveled in the first 30 minutes after the cocaine injection.

To investigate the impact of repeated handling and injection procedure on locomotor responses in the absence of cocaine, one group of male and female mice underwent EcoHIV or sham infection (n = 6/group) and underwent identical handling and exposure to locomotor chambers, except that mice received only saline injections to serve as cocaine-naïve controls.

### Immunohistochemical assessment of astrocyte markers

At least 48 hours after the cocaine challenge test, mice received an injection of either cocaine (10 mg/kg) or saline 90 min before transcardially perfusion. Mice were then perfused with 4% paraformaldehyde (PFA) in 1X PBS and fixed brains were isolated and sectioned (40μm thickness) into triplicate wells for immunohistochemistry. To determine the effect of EcoHIV infection on NAc astrocytes, the NAc brain sections were prepared for immunohistochemical staining of astrocyte markers, Sox9 and glial fibrillary acidic protein (GFAP). For Sox-9, sections were blocked using 10% normal donkey serum (NDS) for 1 hour, then incubated in primary (goat anti-Sox9, 1:200, R&D Systems, AF3075) with 5% NDS overnight. After washing, sections were incubated in the dark in Alexa Fluor® 488 donkey anti-goat secondary (1:250, Jackson ImmunoResearch, 705-545-147) for 2 hours incubation. Tissues were mounted on plus slides and coverslipped with Prolong antifade mountant (Thermo Fisher, P36970). Photomicrographs of 3 sections of 9 fluorescence images in the NAc (bregma: AP: +0.98mm, +1.21mm, +1.42mm) were obtained by fluorescence microscope (Nikon, 10x) and imported into ImageJ, the counts per area (total counts in the selected area, measured as counts/mm^2^) and average intensity per Sox9 (total pixels in selected area normalized by total counts) were analyzed.

Another portion of brain sections were incubated overnight in primary rabbit anti-GFAP (1:10,000, Millipore Sigma, G9269). On the second day, sections were incubated for 30 min with a biotinylated Donkey anti-rabbit antibody (1:1000, Jackson ImmunoResearch, 711-065-152), then transferred to an avidin–biotin complex for 1 h (Vector Laboratories, PK-6200) and developed using diaminobenzidine (DAB, Vector Laboratories, SK-4100). Tissues were mounted on plus slides and coverslipped in DPX mounting media (Electron Microscopy Sciences, 13512). Images were obtained by using a light microscope (Nikon, 10x). Three sections of NAc were imaged (bregma: AP: +0.98mm, +1.21mm, +1.42mm) and analyzed using ImageJ to calculate the percentage per area stained (% pixel in selected area).

### Chemogenetic activation of astrocyte Gq-signaling

A separate cohort of mice was used to determine the effect of chemogenetic activation of astrocytes on cocaine sensitization. Male (n = 16) and female (n = 16) mice were bilaterally injected into the nucleus accumbens (NAc) (bregma: AP +1.3 mm, ML ±1.0 mm, DV –4.8 mm) with either AAV-Gfap-hM3Dq(Gq)-mCherry (0.2 μL per side, 0.1μL/min, gift from Bryan Roth; Addgene viral prep # 50478-AAV5; http://n2t.net/addgene:50478; RRID: Addgene_50478) or control virus (AAV-Gfap-mCherry; 0.2 μL per side, Addgene viral prep #58909-AAV5; http://n2t.net/addgene:58909; RRID:Addgene_58909). Mice recovered for 7 days then underwent EcoHIV inoculation and 5 daily cocaine injections as described above. The only difference in testing was on the challenge test in which, after 60 min of habituation in the open field chamber, mice received an injection of CNO rather than saline (2 mg/kg i.p., n = 16 control virus- or DREADD virus-injected) prior to recording activity for 60 min.

### Immunohistochemical confirmation of AAV placement

To verify virus expression in the NAc, 40 μm brain slices from control virus- or DREADD virus-expressing mice underwent RFP (for mCherry tag) immunofluorescent staining. Slices were incubated in chicken anti-RFP primary (1:1,000, Novus Biologicals, Centennial, CO, NBP1-97371) overnight, then Alexa Fluor® 594 donkey anti-chicken secondary (1:250, Jackson ImmunoResearch, 703-585-155) for two hours. Using fluorescent microscopy, we confirmed that only mice with RFP signal in the NAcore/shell region between 0.98mm and 1.70mm anterior to bregma were included. Distinctions between core and shell were not investigated, but RFP signal was observed in both core and shell. RFP positive signals were confirmed in all surgerized mice. Three mice were excluded due to incorrect placement or insufficient DREADD expression.

To confirm the effect of CNO on Gq signaling, all mice that underwent surgery were subjected one CNO injection (2mg/kg, i.p.) and perfused 90 min later for cFos induction in the NAc. Images of NAc (bregma: AP: +0.98mm, +1.21mm, +1.42mm) in mice with control virus- or DREADD virus were acquired and analyzed for cFos expression (counts/mm^2^).

### Statistical Analyses

GraphPad Prism (10) was used for statistical analysis of behavioral and molecular data. Data were analyzed using unpaired *t*-test or two-way ANOVA. Significant interactions were followed using Holm-Šídák’s multiple comparisons *post hoc* analysis or Fisher’s LSD multiple comparisons test where appropriate. Correlational analyses were performed using simple linear regression. Significance levels for each test were at p < 0.05. Principal component analysis (PCA) was applied to identify common characteristics among the movement variables using R Studio. Raw expression data of cytokine/chemokine were scaled and centered. The correlation matrix and eigenvalue were calculated, eigenvalues greater than 1 were considered significant in contributing to the components, and data were plotted on the first two components. A z-score two population comparison was applied to compare the proportion of mice assigned to each quadrant following PCA.

## Results

### EcoHIV did not impact acute and repeated cocaine-induced locomotor activity

To assess whether EcoHIV infection impacted cocaine-induced locomotion and locomotor sensitization, we analyzed the cocaine-induced locomotor responses of sham and EcoHIV-infected mice across the initiation and expression of locomotor sensitization (see **Figure 1A for timeline**). Daily locomotor activity was analyzed following cocaine exposure using two-way ANOVA. Across the 5 daily cocaine injections, distance traveled was increased in both sham and EcoHIV-infected mice in response to the repeated cocaine injection [two-way ANOVA, main effect of cocaine: F(2.756, 90.93) = 3.198, p = 0.0307; Greenhouse-Geisser corrected; **Figure 1B**]. No main effects of EcoHIV [F(1, 33) = 2.146, p = 0.1524] or EcoHIV by cocaine interaction [F(4, 132) = 0.3758, p = 0.8256] were observed on distance traveled. To determine if the time course of locomotor activity was impacted by EcoHIV infection, we analyzed the change in distance traveled during the 60 min open field sessions in 5-min bins on the first (day 1) and final (day 5) session of daily cocaine exposure. On day 1, both sham and EcoHIV-infected mice increased activity following cocaine injection, but no main effect of EcoHIV or interaction were observed [two-way ANOVA; main effect of time: F(1.239, 39.63) = 16.87, p <0.0001; interaction: F(11, 352) = 0.07215, p > 0.9999, main effect of EcoHIV: F (1, 32) = 0.01614, p = 0.8997, **Figure 1C**]. However, on day 5, we observed an EcoHIV and time interaction on distance traveled [two-way ANOVA; interaction: F(11, 363) = 1.844, p = 0.0456; main effect of time: F(1.260, 41.59) = 18.26, p < 0.0001; no main effect of EcoHIV: F(1, 33) = 1.878, p = 0.1798, Greenhouse-Geisser corrected, **Figure 1D**]. Post hoc Holm-Šídák’s multiple comparisons test revealed that the change in distance traveled was not significant between sham and EcoHIV-infected mice at any time point. This finding suggested that EcoHIV-infected mice showed a trend toward higher cocaine locomotor responses after 5 daily exposures to cocaine.

### EcoHIV significantly increased cocaine locomotor sensitization on the challenge test

After induction of cocaine-induced locomotor response by repeated injection, we next investigated whether EcoHIV infection impacted the expression of cocaine-induced locomotor sensitization after a 6-day cocaine-free period. We observed greater cocaine-induced increases in distance traveled in EcoHIV-infected mice than in sham control mice [t(33) = 2.257, p = 0.0308, unpaired t-test; **Figure 2B**]. To investigate the time course of this effect, change in activity was analyzed across 5-min bins (**Figure 2A**). A main effect of time after cocaine injection [F(1.241, 40.96) = 114.4, p < 0.0001] and a main effect of EcoHIV [F(1, 33) = 4.479, p = 0.042] were observed. Further, EcoHIV and time interacted to impact distance traveled [two-way ANOVA; interaction: F(11, 363) = 2.398, p = 0.007]. A post hoc Holm-Šídák’s multiple comparisons test revealed that the change in distance traveled in EcoHIV mice was greater than sham controls from 20 - 45 min after cocaine administration (p’s < 0.05).

**Figure 2.**
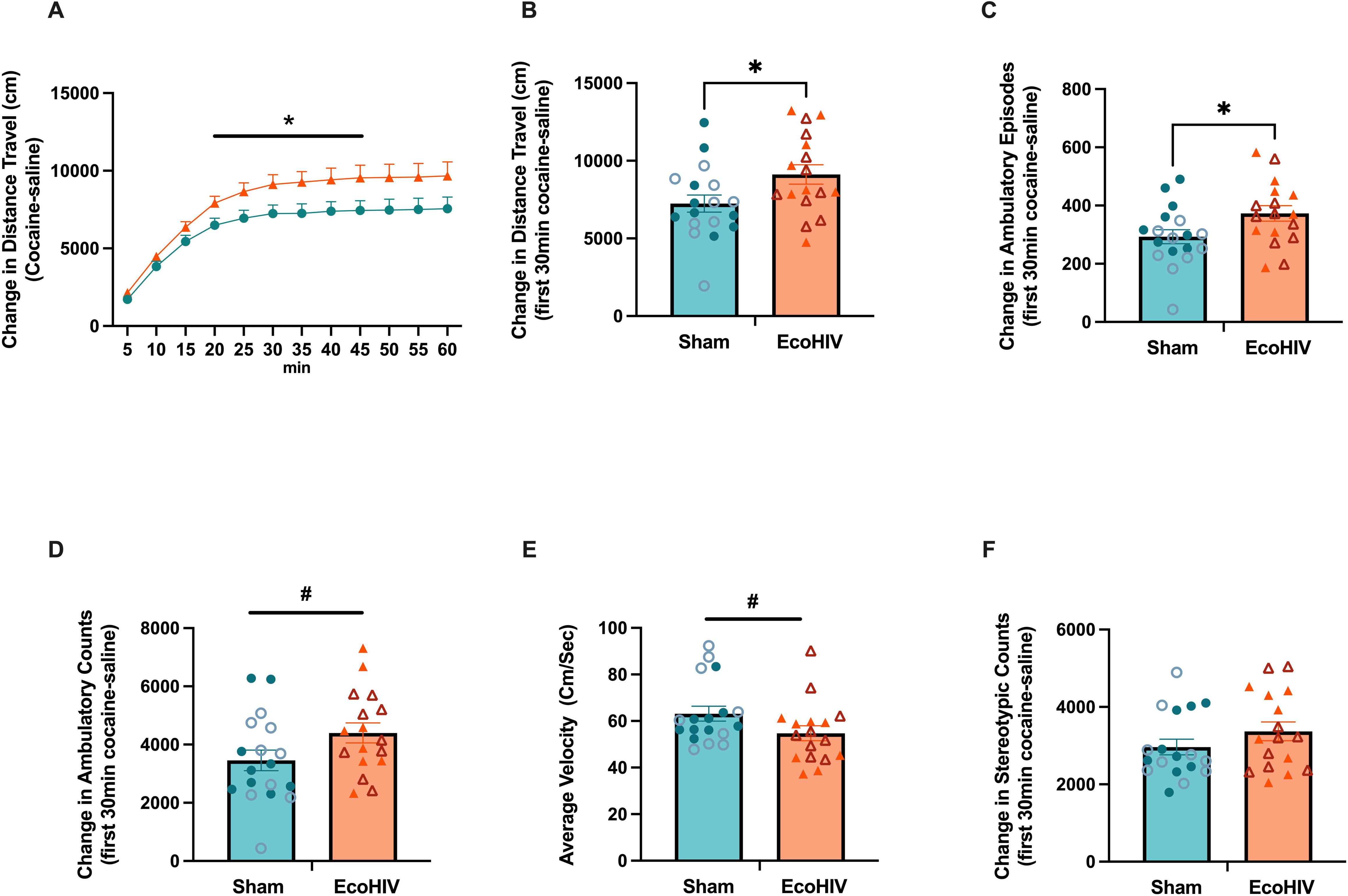
Expression of cocaine locomotor sensitization in the challenge test. EcoHIV significantly increased the locomotor response to cocaine, as measured by an increase in distance traveled across the session (**A**) and in total (**B**) when compared to sham mice. (**C**) EcoHIV-infected mice increased ambulatory episodes to a greater degree than sham mice. EcoHIV-infected mice exhibited a trend toward higher ambulatory counts (**D**) and lower average velocity (**E**). No difference was observed between EcoHIV and sham mice in stereotypy counts (**F**). Circle and triangle symbols represent sham and EcoHIV mice; Closed and open symbols represent male and female mice, respectively. n = 17 - 18/group. Bars represent means +/- SEM. # p < 0.1, * p < 0.05.

### EcoHIV increased cocaine-induced activity

To further characterize the effect of EcoHIV infection on components of locomotor sensitization, we investigated multiple parameters including ambulatory counts, ambulatory episodes, average velocity, and stereotypy counts. EcoHIV-infected mice exhibited a higher number of ambulatory episodes [t(33) = 2.231, p = 0.0326; unpaired t-test; **Figure 2C**] and showed a trend toward a higher number of ambulatory counts in response to the cocaine challenge compared to sham controls [t(33) = 1.912, p = 0.0646; unpaired t-test; **Figure 2D**]. Interestingly, EcoHIV-infected mice displayed a trend toward reduced velocity during the challenge test [t(33) = 1.845, p = 0.0741; unpaired t-test; **Figure 2E**]. Both EcoHIV-infected mice and sham controls showed similar stereotypic behavior to the cocaine challenge [t(33) = 1.289, p = 0.2063; unpaired t-test; **Figure 2F**]. These findings suggest that EcoHIV affects cocaine-induced locomotor sensitization in ways that go beyond merely increasing all types of movement.

To further characterize locomotor changes in response to a cocaine challenge, we applied principal component analysis (PCA), a dimension reduction approach, to identify differences in performance between EcoHIV-infected and sham controls using distance traveled, ambulatory counts, ambulatory episodes, average velocity, and stereotypy counts (**Figure 3A**). The PCA identified two components that captured the majority of the variance (87.3%). Distribution along the first principal component (PC1) was driven by distance traveled, ambulatory counts, ambulatory episodes, and velocity, explaining 69.9% of the cumulative variance. The velocity opposed the other three movement-related variables, indicating velocity was negatively correlated with other movement metrics which clustered together. The second principal component (PC2) was driven by stereotypy counts, accounting for 17.4% of the variance. This suggests that stereotypy was largely independent of movement and velocity in the cocaine challenge.

**Figure 3.**
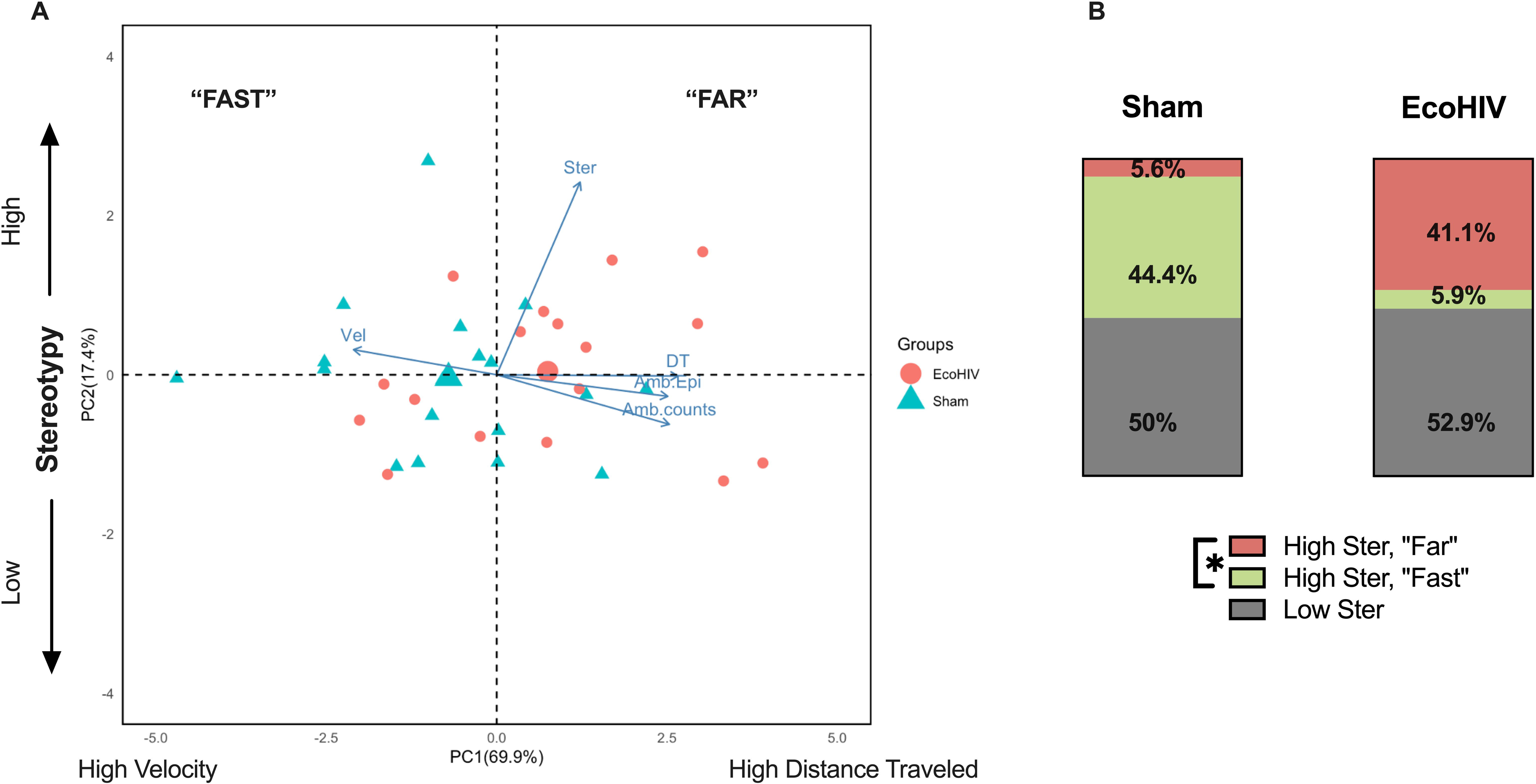
Principal component analysis (PCA) of locomotor changes in response to a cocaine challenge. (**A**) Distribution along the first principal component (PC1) was driven by distance traveled, ambulatory counts, ambulatory episodes, and velocity, explaining 69.9% of the cumulative variance. The second principal component (PC2) accounted for 17.4% of the variance and was positively correlated with stereotypy counts. (**B**) The cluster profile of EcoHIV-infected and sham control showed similar distribution along PC2, with ∼50% in the high stereotypy cluster. Within high stereotypy cluster, EcoHIV-infected mice were more likely to exhibit "Far" behavior (41.1%) compared to sham mice (5.6%), while sham mice were more likely to exhibit "Fast" behavior (44.4%) compared to EcoHIV-infected mice (5.9%). This indicates that EcoHIV infection shifts cocaine-induced locomotor activity towards slower but sustained movements during the challenge. *p < 0.05.

We found that sham and EcoHIV-infected mice were similarly distributed along PC2 (high stereotypy vs. low stereotypy, Sham: 50% high, EcoHIV: 52.9% high), with ∼50% of each group in the high stereotypy cluster. However, among the high stereotypy population, EcoHIV and sham mice differed along PC1. Among mice with high stereotypy, 41.1% of EcoHIV mice were in the high movement and low velocity cluster (defined as "Far"), and 5.9% were in the low movement and high velocity cluster (defined as "Fast"). In contrast, the sham mice exhibited a different distribution: 5.6% were in the "Far" cluster, and 44.4% were in the "Fast" cluster (z-score two population comparison; z = -2.5083, p = 0.01208; **Figure 3B**). These results indicate that EcoHIV infection selectively shifted cocaine-induced locomotor activities toward slower, but longer-lasting movements during the challenge.

### EcoHIV infection increased the number of Sox9 positive cells in the NAcore

To determine if the increased cocaine sensitization observed in EcoHIV-infected mice was associated with alterations in astrocytes in the NAc, we performed immunohistochemistry analysis and quantified the expression of Sox9, a marker specific to reactive astrocytes in adult mice, and expression of glial fibrillary acidic protein (GFAP) in the NAc. Sox9 counts per area (mm²) and average intensity in the NAcore and NAshell were compared between sham and EcoHIV-infected animals. Analysis of Sox9+ cells per area revealed that EcoHIV infection increased the number of Sox9+ cells in the NAcore [t(22) = 3.035, p = 0.0061, unpaired t-test, **Figure 4A**], but not in the NAshell [t(22) = 0.3523, p = 0.7280, unpaired t-test, **Figure 4B**]. The average immunofluorescence intensity of Sox9 was also analyzed to determine the average expression level per cell. No effect of EcoHIV infection on Sox9 intensity was observed in the NAcore [t(22) = 1.121, p = 0.2744, **Figure 4C**] or in the NAshell [t(22) = 0.8986, p = 0.3786, **Figure 4D**]. These findings indicate that EcoHIV infection increased the number of Sox9+ cells in the NAcore rather than the expression level of Sox9 in each astrocyte. Representative images of Sox9 expression in the NAcore and NAshell of EcoHIV-infected and sham controls are shown in **Figure 4E**.

**Figure 4.**
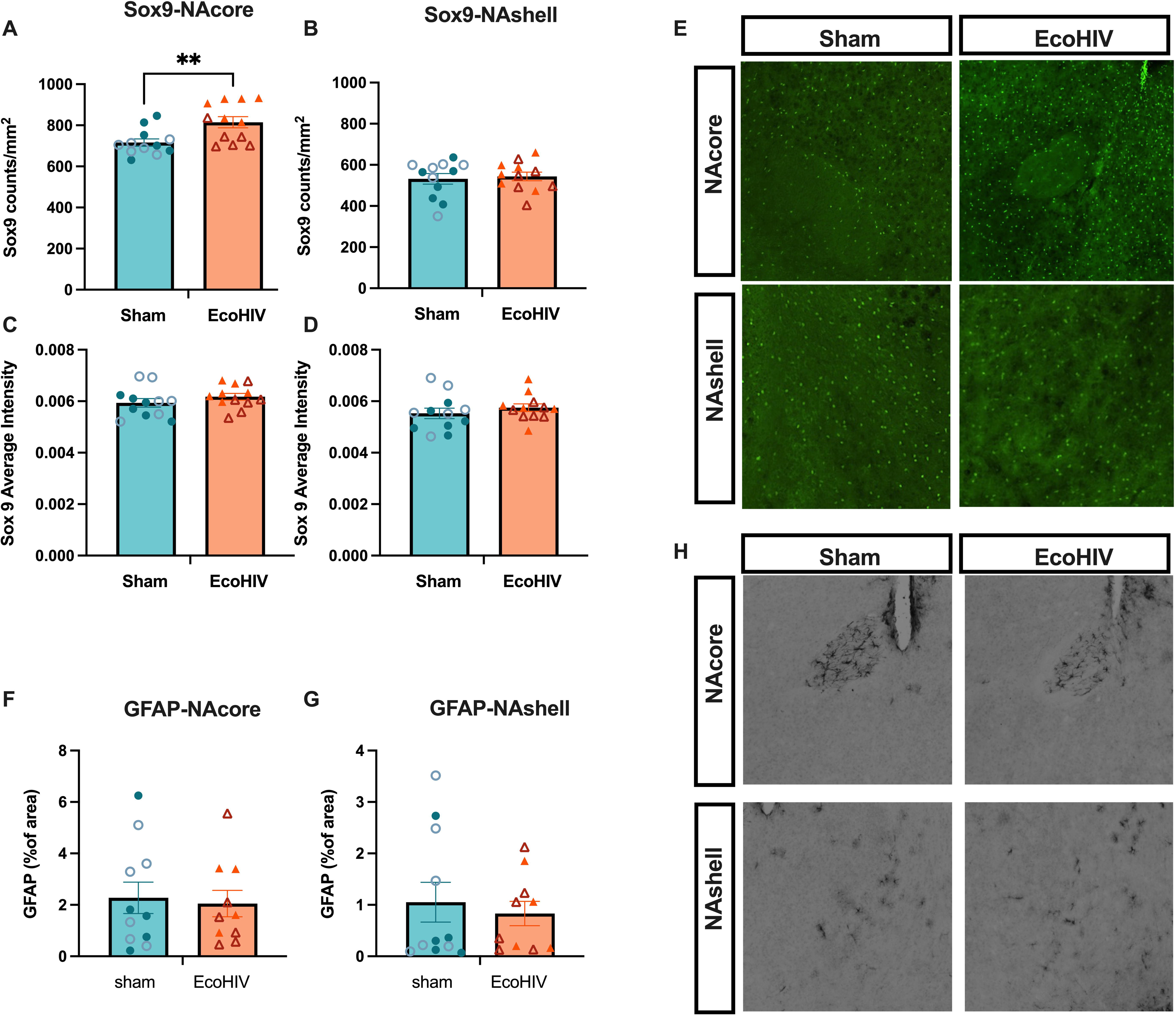
Expression of astrocyte markers in subregions of the NAc. EcoHIV infection increased the number of Sox9 positive cells (counts/mm2) in the NAcore (**A**) but not NAshell (**B**). No effect of EcoHIV infection on Sox9 intensity was observed in the NAcore (**C**) or NAshell (**D**). (**E**) Representative immunofluorescent images of Sox9 expression in the NAcore and NAshell in sham or EcoHIV-infected mice. No significant differences in the % of area of GFAP expression between EcoHIV-infected and sham mice in both the NAcore (**F**) and NAshell (**G**) were observed. (**H**) Representative immunohistochemistry images of GFAP in the NAcore and NAshell. Closed and open symbols represent male and female mice, respectively. Data represent mean +/- SEM. *p < 0.05.

The percentage per area (% of area) of GFAP protein in the NAc was compared between EcoHIV-infected and sham mice. An unpaired t-test revealed no significant difference in the % of area between EcoHIV-infected and sham mice in both the NAcore [t(19) = 0.2767, p = 0.7850, **Figure 4F**] and NAshell [t(19) = 0.4748, p = 0.6404, **Figure 4G**]. Representative images of GFAP expression in the NAcore and NAshell of EcoHIV-infected and sham controls are shown in **Figure 4H**.

### Chemogenetic activation of Gq signaling in NAc astrocytes suppressed cocaine-induced hyperlocomotion in EcoHIV-infected mice

To determine whether Gq signaling in NAc astrocytes could modulate the expression of cocaine sensitization in EcoHIV-infected mice, locomotor response was assessed following chemogenetic activation of Gq signaling in mice expressing the Gq-DREADD (or control virus) under the GFAP promoter. **Figure 5A**). We found that EcoHIV infection and DREADD condition interacted to affect distance travel [F (1, 30) = 5.868, p = 0.0217; Two-way ANOVA], but no main effects of DREADD or of EcoHIV were detected [main effect of DREADD: F (1, 30) = 0.4396, p = 0.5124; main effect of EcoHIV: F (1, 30) = 0.08449, p = 0.7733]. A post hoc Fisher’s LSD multiple comparisons test revealed that in mice expressing control virus (non-DREADD), EcoHIV mice showed a trend towards higher distance travel compared to sham mice (p = 0.0646). However, in DEADD-expressing mice, CNO administration significantly attenuated cocaine sensitization in EcoHIV-infected mice (p = 0.0424, **Figure 5B**). This suggests that the hypersensitization induced by EcoHIV could be reproduced in surgerized mice, and that chemogenetic activation of astrocyte signaling in the NAc can reverse EcoHIV-induced hypersensitization to cocaine.

**Figure 5.**
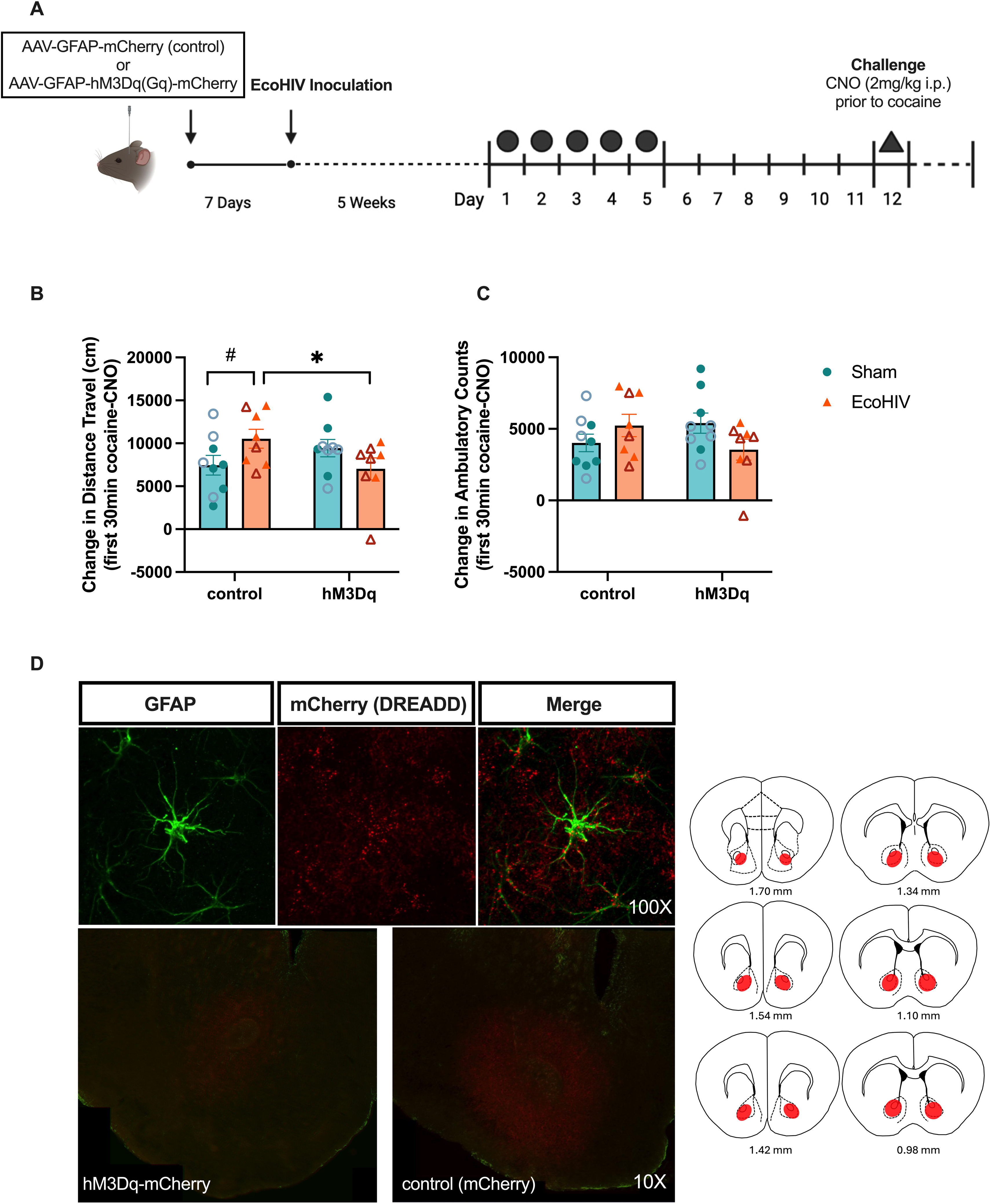
Chemogenetic activation of Gq signaling in NAc astrocytes. (**A**) Timeline of chemogenetics study. Astrocytic Gq-DREADD (AAV-Gfap-hM3Dq(Gq)-mCherry) or control (AAV-Gfap-mCherry) virus was injected to the NAc of mice prior to EcoHIV inoculation. Five weeks after inoculation, mice underwent the cocaine sensitization test. On the challenge day, mice received an injection of CNO to activate astrocyte Gq signaling in the NAc before cocaine challenge. (**B**) Astrocyte Gq-DREADD activation by CNO administration significantly attenuated distance traveled in EcoHIV-infected mice. (**C**) Ambulatory counts were not significantly reduced following Gq-DREADD activation. (**D**) Colocalized GFAP and RFP (hM3Dq-mCherry) indicated selective DREADD expression in astrocytes. Representative immunofluorescence images of hM3Dq-mCherry or mCherry in the NAc sections. Data represent mean +/- SEM. Closed and open symbols represent male and female mice, respectively. n= 8-9 /group. #p < 0.1, *p < 0.05.

A two-way ANOVA revealed a significant interaction between DREADD and EcoHIV on ambulatory counts [F (1, 30) = 4.706, p = 0.0381], but no main effect of DREADD [F (1, 30) = 0.04562, p = 0.8323] or EcoHIV [F (1, 30) = 0.2010, p = 0.6571], were observed (**Figure 5C**). However, Fisher’s LSD multiple comparisons post hoc analyses showed no significant comparisons (p’s > 0.05). There was no difference between DREADD and EcoHIV for ambulatory episodes [interaction: F (1, 30) = 2.162. p = 0.1518; main effect of DREADD: F (1, 30) = 0.2595, p = 0.6142; main effect of EcoHIV: F (1, 30) = 0.05751, p = 0.8121], average velocity [interaction: F (1, 30) = 0.2223, p = 0.6407; main effect of DREADD: F (1, 30) = 0.1105, p = 0.7419; main effect of EcoHIV: F (1, 30) = 0.03418, p = 0.8546], and stereotypy counts [interaction: F (1, 30) = 1.058, p = 0.3120; main effect of DREADD: F (1, 30) = 0.2953, p = 0.5909; main effect of EcoHIV: F (1, 30) = 1.008, p = 0.3234]. Representative immunofluorescence images of hM3Dq-mCherry or mCherry in the NAc sections (**Figure 5D**). Colocalized GFAP and RFP (hM3Dq-mCherry) indicated selective DREADD expression in astrocytes.

### EcoHIV-NDK spleen DNA load correlated with expression of cocaine sensitization

To validate the infection status of mice, we isolated and purified DNA from the spleens of all EcoHIV-infected mice and representative sham mice, and quantified HIV-1 long terminal repeat (LTR) DNA levels via qPCR. All EcoHIV-infected mice showed detectable levels of viral DNA. No viral DNA was detected in sham control mice. We verified that EcoHIV LTR DNA levels were in the range from 0.3 to 8 × 10^3^ viral DNA copies per 10^6^ spleen cells at the end of experiments (8 weeks following inoculation, **Figure 6A**). Viral DNA levels were compared between cocaine-naïve and cocaine-exposed EcoHIV-infected mice. An unpaired t-test revealed no main effect of cocaine were observed [t(22) = 0.3657, p = 0.7181, unpaired t-test, **Figure 6A**], indicating EcoHIV-NDK splenic viral DNA burden was not impacted by a history of cocaine administration in these conditions.

**Figure 6.**
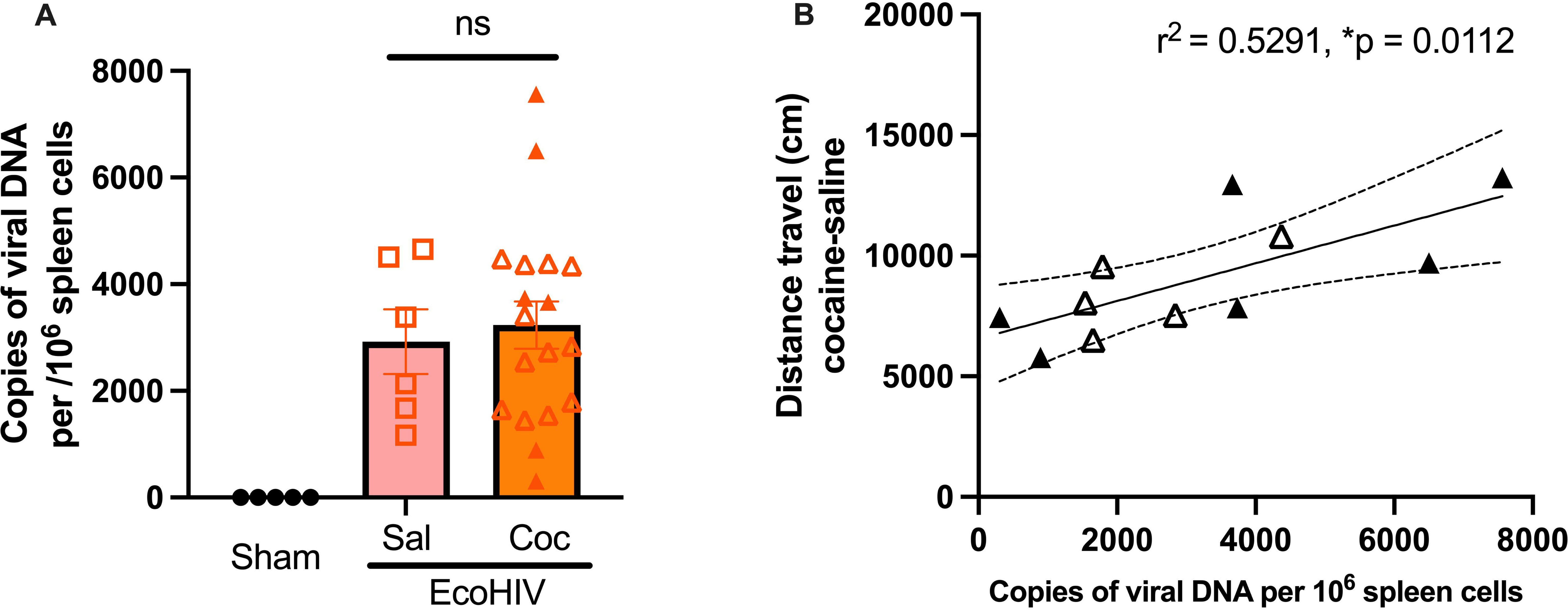
EcoHIV-NDK spleen DNA load correlated with expression of cocaine sensitization. (**A**) Copies of EcoHIV-NDK spleen viral DNA at the end of experiment (week 8 of infection). The number of copies of viral DNA per 10^6^ spleen cells did not differ between cocaine-naïve and cocaine exposed EcoHIV-infected mice. (**B**) Linear regression showed a positive correlation between the DNA viral load and distance travel. Closed and open symbols represent non-surgerized and surgerized mice, respectively. DREADD-expressing mice were excluded from the regression analysis. *p < 0.05.

To determine the relationship between EcoHIV viral DNA level and the expression of cocaine sensitization, we performed simple linear regression analysis between viral DNA level and distance traveled in EcoHIV-infected mice (EcoHIV-infected mice expressing DREADD were excluded). Linear regression showed a positive correlation between the DNA viral load and distance travel (R^2^= 0.5291, p = 0.0112, **Figure 6B**). This result may indicate the degree of cocaine sensitization is associated with EcoHIV DNA level.

## Discussion

Our findings demonstrate EcoHIV infection facilitated behavioral sensitization in response to cocaine and altered NAc astrocytes, and further that behavioral alterations could be rescued through chemogenetic activation of Gq signaling in NAc astrocytes. In particular, we observed that EcoHIV infection selectively promoted the expression of cocaine locomotor sensitization in response to cocaine challenge, without impacting initial locomotor response to cocaine. This was characterized by EcoHIV infection-driven shifts in cocaine-induced movement toward a lower velocity but higher distance pattern of activity during challenge test. As astrocytes in the NAc are disrupted by cocaine exposure, and modulating astrocytes can attenuate cocaine-seeking behavior, we investigated whether astrocytes were altered by EcoHIV in this model. We found that EcoHIV infection was associated with increased expression of Sox9 positive cells – a potential marker of reactive astrocytes in the NAcore, but not NAshell, in the cocaine sensitized mice. We further demonstrated that increasing astrocytic Gq signaling through chemogenetics could attenuate EcoHIV-induced cocaine sensitization.

Psychostimulant-induced behavioral sensitization is particularly interesting because it not only serves as a model for the pathophysiological changes resulting from repeated administration of cocaine but also reflects the neuroadaptations and behavioral plasticity contribute to cocaine use disorder (Thomas et al., 2001; Li et al., 2004; Steketee and Kalivas, 2011). It has been proposed there are two distinct phases of behavioral sensitization to cocaine: initiation and expression. The initiation phase involves acute pharmacological effects of cocaine and transient changes in neurotransmission, leading to enduring neuroplastic alterations that facilitate the expression of sensitization (Kalivas and Duffy, 1990; Riday et al., 2012). Our data revealed that sham and EcoHIV-infected mice did not differ in distance traveled after first cocaine injection (acute cocaine exposure) following 5 weeks of EcoHIV infection (**Figure 1C**). This was unexpected as acute cocaine-induced locomotor responses are dependent on innate dopaminergic function (Kalivas and Duffy, 1990), and others have observed impairments in dopaminergic plasticity that are implicated in neuroHIV and contribute to reward deficits in both EcoHIV and other HIV models (Maragos et al., 2002; Kesby et al., 2016; Gaskill et al., 2017; Olson et al., 2018), Five weeks of EcoHIV infection alone may not affect dopamine release by acute cocaine injection or dopamine transporter function in regulating extracellular dopamine level. However, we revealed a trend towards a greater response in EcoHIV-infected mice after 5 days of cocaine exposure, suggesting that neuroadaptations that facilitate the development of locomotor sensitization may occur more rapidly in the EcoHIV exposed animals (**Figure 1D**).

Cocaine motor response continues to increase after cessation of cocaine use, which promotes long-lasting sensitization refers to expression of sensitization (Van Zessen et al., 2021). In our study, we observed a prolonged locomotor response across the session in EcoHIV-infected mice compared to sham controls (**Figure 2A**). Additionally, our PCA analysis indicates that EcoHIV infection shifted the locomotor profile toward slower but more sustained movements during the challenge test (**Figure 3B**). This hyper-responsiveness following the cessation of cocaine is associated, in part, with dopaminergic neuroadaptations in the NAc that facilitate the expression of cocaine locomotor sensitization (Kalivas and Duffy, 1990; Boudreau et al., 2007; Ghasemzadeh et al., 2009; Harraz et al., 2021; Van Zessen et al., 2021). Repeated cocaine exposure reduces the availability of the dopamine transporter (DAT) in the NAc, leading to an increased concentration of dopamine in response to subsequent cocaine challenges (Kalivas and Duffy, 1990; Harraz et al., 2021). Moreover, previous research has shown that increased locomotor sensitization in response to psychostimulants in HIV rodent models occurs due to alterations in dopamine function. For example, intracranial exposure to the HIV protein Tat enhances the locomotor sensitization induced by both cocaine and methamphetamine (Maragos et al., 2002; Harrod et al., 2008; Liu et al., 2009; Paris et al., 2014, 2015; Kesby et al., 2017; Basova et al., 2020). Disruption of the dopamine system by HIV and its viral proteins, such as through dysregulation of DAT, can promote hyperdopaminergic tone within the striatum of HIV rodent models (Cass et al., 2003; Ferris et al., 2010; Nickoloff-Bybel et al., 2020). These findings align with our results, suggesting that EcoHIV infection, in combination with repeated cocaine exposure, may synergistically enhance cocaine responsiveness and lead to more pronounced sensitization effects (Liu et al., 2009; Ferris et al., 2010; Kesby et al., 2017).

The mechanisms underlying the neurobehavioral outcomes of repeated cocaine exposure, and the modulatory effect of HIV, also involve alterations in glutamatergic plasticity. For example, repeated cocaine exposure and cocaine withdrawal increase glutamate receptor (e.g. AMPA and NMDA) trafficking to synaptic membranes and enhances glutamate transmission in the NAc (Ghasemzadeh et al., 2009). These adaptions prime LTP in the NAc and promote the expression of cocaine-induced locomotor sensitization as well as reward motivation and memory (Kalivas, 2004; Boudreau et al., 2007; Huang et al., 2009; Brown et al., 2011). HIV and its viral proteins can compromise glutamatergic system signaling to exacerbate cocaine-related behavioral outcomes. For example, gp120 and Tat induce deficits in the corticostriatal glutamate system, leading to synaptic degeneration and dysfunction in the projections from the prefrontal cortex (PFC) to the NAc, which are primarily glutamatergic and mediate flexible cocaine-seeking and relapse behaviors (Wang et al., 2003; Gupta et al., 2010; Paris et al., 2014; Du Plessis et al., 2015; Kesby et al., 2016; Melendez et al., 2016). Specifically, gp120 and Tat directly activate glutamate receptors that contribute to synaptic overactivation and synaptic damage (Eugenin et al., 2007; McLaurin et al., 2018; Gorska and Eugenin, 2020). Moreover, parallel to the current sensitization study, we have previously demonstrated that cocaine-seeking behavior is positively correlated with NMDA receptor (GluN2A) levels in the nucleus accumbens (NAc) of EcoHIV-infected mice (Namba et al., 2024). These findings suggest that EcoHIV infection can affect glutamatergic signaling in the reward pathway, a disruption that is further exacerbated by cocaine, leading to alterations in locomotor sensitization.

Astrocytes are an important regulator of glutamate uptake and clearance, a key mechanisms underlying motivated cocaine seeking (Kalivas, 2009; Knackstedt et al., 2010; Reissner et al., 2015). Cocaine exposure and withdrawal impair the glutamate transporter (GLT-1) and cystine-glutamate antiporter (Sxc-) within the NAc, leading to increased extracellular levels of glutamate and postsynaptic excitability within the NAc following exposure to cocaine or cocaine-associated cues (McFarland et al., 2003; Kau et al., 2008; Reissner et al., 2015; Kim et al., 2018). Astrocytes are also affected by HIV. HIV infects astrocytes and induces astrocyte reactivity that triggers release of astrocyte-mediated immune factors (Nooka and Ghorpade, 2017) and reduction of GLT-1 expression and associated glutamate transport (Wang et al., 2003; Melendez et al., 2016). Our data reveal that EcoHIV infection increased the number of astrocytes expressing Sox9 – a potential marker of reactive astrocytes - in the NAcore of cocaine-exposed mice (Sun et al., 2017). Further, our previous findings revealed that 5 weeks of EcoHIV infection alone increases pro-inflammatory cytokines and microglia reactivity in the NAc (Xie et al., 2024). Microglia can release these pro-inflammatory cytokines in response to HIV insult, which in turn activates astrocytes (Hong et al., 2022; Namba et al., 2023), consistent with our finding that EcoHIV increased astrocyte reactivity in the NAc. These findings may indicate NAcore astrocytes are susceptible to both the independent and interactive effects of EcoHIV and repeated cocaine exposure. Prolonged exposure to pro-inflammatory environment can cause astrocyte activation to become pathological (Pittaluga, 2017). Thus, reactive astrocytes become less efficient in glutamate regulation and release more cytokines which could accelerate neuronal adaptations that drive locomotor sensitization in response to a subsequent cocaine challenge.

Neuronal plasticity requires communication between neurons and astrocytes (Allen and Eroglu, 2017). Astrocyte Ca^2+^ signaling promotes the release of substrates such as glutamate, GABA, and ATP/adenosine, which can interact with neuronal receptors and thus shape neuronal plasticity (Goenaga et al., 2023). For example, astrocyte Ca^2+^ events regulate the release of extrasynaptic glutamate, stimulating presynaptic metabotropic glutamate group II receptors (mGlu2/3) to suppress excess drug- and drug cue-evoked synaptic glutamate release (Koob and Volkow, 2010; Scofield et al., 2015). Additionally, dopamine activates astrocyte Ca^2+^ activity in the nucleus accumbens (NAc) through astrocytic D1 receptors. This astrocyte Ca^2+^ signaling stimulates ATP/adenosine release, which depresses excitatory neuronal transmission via the activation of presynaptic A1 receptors (Corkrum et al., 2020). Cocaine has been shown to induce a series of Ca^2+^-associated responses in the NAc astrocytes that influence the cocaine behavioral outcomes (Corkrum et al., 2020; Wang et al., 2021a, 2021b). Chemogenetic activation of astrocyte Ca^2+^ signaling via Gq-DREADD in the NAc prevents cue-induced cocaine and methamphetamine seeking (Scofield et al., 2015; Siemsen et al., 2019), potentially through the restoration of glutamate homeostasis via the mGlu2/3 signaling in the NAc. HIV infection can also have profound effects on dysregulation of astrocytic Ca^2+^ signaling. For instance, HIV Tat protein upregulates astrocyte reactivity and induces astrocyte-mediated neurodegeneration in a calcium-dependent manner (Pandey et al., 2022). Our results demonstrated that stimulation of astrocytic Gq signaling in the NAc was able to effectively suppress hypersensitization to cocaine in EcoHIV-infected mice (**Figure 5B**). This may suggest that EcoHIV infection is associated with dysregulation of astrocyte Ca^2+^ activity to facilitate the expression of cocaine-induced locomotor sensitization. It also reveals astrocytes as a potential therapeutic target for comorbid HIV and cocaine use disorder.

One key finding from the present study is that the distance traveled following a cocaine challenge positively correlated with HIV LTR DNA levels in the spleens of EcoHIV-infected mice. Despite little to no viral load at the time of testing (Gu et al., 2018), full-length, linear, unintegrated viral DNA is abundant in brain and lymphoid tissue during the asymptomatic phase of infection (Pang et al., 1990; Chun et al., 1997; Teo et al., 1997). Moreover, viral proteins such as Tat and Nef can be synthesized from unintegrated viral DNA (Wu et al., 2004), which represents a potential mechanism through which elevated viral DNA burden may contribute to enhanced sensitivity to the psychomotor effects of cocaine. Indeed, HIV-1 Tat transgenic mice exhibit potentiated cocaine-induced locomotor activity (Paris et al., 2014), and HIV-1 transgenic rats (which constitutively express Tat, Nef, among other viral proteins) exhibit an enhanced sensitivity to cocaine evidenced by a leftward shift in the cocaine dose response function (McIntosh et al., 2015). Our group has also demonstrated enhanced cocaine-induced locomotor activity following daily, repeated exposure to cocaine within a cocaine conditioned place preference paradigm (Namba et al., 2024). While speculative, it is certainly possible that enhanced transcriptional activity of viral proteins such as Tat or Nef could contribute to increased cocaine-induced locomotor activity among EcoHIV-infected mice despite low/undetectable viral load. Future studies quantifying both systemic and CNS expression levels of these viral substrates would be informative.

## Conclusion

Here, we demonstrate that EcoHIV infection increased locomotor sensitization to cocaine, which was further associated with astrocytic adaptations within the NAc. Chemogenetic activation of NAc astrocyte signaling was able to reverse EcoHIV-induced hyper-sensitization, highlighting the potential of astrocytes as a therapeutic target to mitigate cocaine-related behaviors among people living with HIV. These insights could inform new therapeutic strategies for managing cocaine use disorder in the context of HIV infection.

## Acknowledgements

This research was supported by NIH award DP2DA051907 (JMB), R21DA056309 (JGJ), R01AG081929 (JGJ), and pilot awards from The Comprehensive Neuro-AIDS Center Grant P30MH092177-9 (JMB and MDN). The EcoHIV-NDK plasmid was a gift from David Volsky.

## References

Allain F, Roberts DCS, Lévesque D, Samaha AN (2017) Intermittent intake of rapid cocaine injections promotes robust psychomotor sensitization, increased incentive motivation for the drug and mGlu2/3 receptor dysregulation. Neuropharmacology 117:227–237.

Allen NJ, Eroglu C (2017) Cell Biology of Astrocyte-Synapse Interactions. Neuron 96:697–708.

Basova L V, Kesby JP, Kaul M, Semenova S, Cecilia M, Marcondes G (2020) Systems Biology Analysis of the Antagonizing Effects of HIV-1 Tat Expression in the Brain over Transcriptional Changes Caused by Methamphetamine Sensitization. Viruses 12:426.

Baum MK, Rafie C, Lai S, Sales S, Page B, Campa A (2009) Crack-cocaine use accelerates HIV disease progression in a cohort of HIV-positive drug users. J Acquir Immune Defic Syndr 50:93–99.

Bazargani N, Attwell D (2016) Astrocyte calcium signaling: The third wave. Nat Neurosci 19:182–189.

Boudreau AC, Reimers JM, Milovanovic M, Wolf ME (2007) Cell Surface AMPA Receptors in the Rat Nucleus Accumbens Increase during Cocaine Withdrawal But Internalize after Cocaine Challenge in Association with Altered Activation of Mitogen-Activated Protein Kinases. J Neurosci 27:10621–10635.

Brown TE, Lee BR, Mu P, Ferguson D, Dietz D, Ohnishi YN, Lin Y, Suska A, Ishikawa M, Huang YH, Shen H, Kalivas PW, Sorg BA, Zukin RS, Nestler EJ, Dong Y, Schlüter OM, Purpura DP (2011) Cellular/Molecular A Silent Synapse-Based Mechanism for Cocaine-Induced Locomotor Sensitization.

Buch S, Yao H, Guo M, Mori T, Su T-P, Wang J (2011) Cocaine and HIV-1 Interplay: Molecular Mechanisms of Action and Addiction. J Neuroimmune Pharmacol 6:503.

Bull C, Freitas KCC, Zou S, Poland RS, Syed WA, Urban DJ, Minter SC, Shelton KL, Hauser KF, Negus SS, Knapp PE, Bowers MS (2014) Rat Nucleus Accumbens Core Astrocytes Modulate Reward and the Motivation to Self-Administer Ethanol after Abstinence. Neuropsychopharmacology 39:2835.

Cass WA, Harned ME, Peters LE, Nath A, Maragos WF (2003) HIV-1 protein Tat potentiation of methamphetamine-induced decreases in evoked overflow of dopamine in the striatum of the rat. Brain Res 984:133–142.

Chun TW, Carruth L, Finzi D, Shen X, DiGiuseppe JA, Taylor H, Hermankova M, Chadwick K, Margolick J, Quinn TC, Kuo YH, Brookmeyer R, Zeiger MA, Barditch-Crovo P, Siliciano RF (1997) Quantification of latent tissue reservoirs and total body viral load in HIV-1 infection. Nat 1997 3876629 387:183–188.

Corkrum M, Covelo A, Lines J, Bellocchio L, Pisansky M, Loke K, Quintana R, Rothwell PE, Lujan R, Marsicano G, Martin ED, Thomas MJ, Kofuji P, Araque A (2020) Dopamine-Evoked Synaptic Regulation in the Nucleus Accumbens Requires Astrocyte Activity. Neuron 105:1036–1047.e5.

Crombag HS, Jedynak JP, Redmond K, Robinson TE, Hope BT (2002) Locomotor sensitization to cocaine is associated with increased Fos expression in the accumbens, but not in the caudate.

Dash S, Balasubramaniam M, Villalta F, Dash C, Pandhare J (2015) Impact of cocaine abuse on HIV pathogenesis. Front Microbiol 6.

Du Plessis S, Vink M, Joska JA, Koutsilieri E, Bagadia A, Stein DJ, Emsley R (2015) HIV infection is associated with impaired striatal function during inhibition with normal cortical functioning on functional MRI. J Int Neuropsychol Soc 21:722–731.

Eugenin EA, King JE, Nath A, Calderon TM, Zukin RS, Bennett MVL, Berman JW (2007) HIV-tat induces formation of an LRP–PSD-95– NMDAR–nNOS complex that promotes apoptosis in neurons and astrocytes. Proc Natl Acad Sci U S A 104:3438.

Ferris MJ, Frederick-Duus D, Fadel J, Mactutus CF, Booze RM (2010) Hyperdopaminergic tone in HIV-1 protein treated rats and cocaine sensitization. J Neurochem 115:885–896.

Gaskill P, Miller D, George J-G, Yano H, Khoshbouei H (2017) HIV, Tat and Dopaminergic Transmission. Neurobiol Dis 105:51.

Ghasemzadeh MB, Vasudevan P, Mueller C (2009) Locomotor sensitization to cocaine is associated with distinct pattern of glutamate receptor trafficking to the postsynaptic density in prefrontal cortex: Early versus late withdrawal effects. Pharmacol Biochem Behav 92:383–392.

Goenaga J, Araque A, Kofuji P, Herrera Moro Chao D (2023) Calcium signaling in astrocytes and gliotransmitter release. Front Synaptic Neurosci 15:1138577.

Gorska AM, Eugenin EA (2020) The Glutamate System as a Crucial Regulator of CNS Toxicity and Survival of HIV Reservoirs. Front Cell Infect Microbiol 10:261.

Gu CJ, Borjabad A, Hadas E, Kelschenbach J, Kim BH, Chao W, Arancio O, Suh J, Polsky B, McMillan JE, Edagwa B, Gendelman HE, Potash MJ, Volsky DJ (2018) EcoHIV infection of mice establishes latent viral reservoirs in T cells and active viral reservoirs in macrophages that are sufficient for induction of neurocognitive impairment. PLoS Pathog 14.

Gupta S, Knight AG, Gupta S, Knapp PE, Hauser KF, Keller JN, Bruce-Keller AJ (2010) HIV-Tat elicits microglial glutamate release: role of NAPDH oxidase and the cystine-glutamate antiporter. Neurosci Lett 485:233–236.

Harraz MM, Guha P, Kang IG, Semenza ER, Malla AP, Song YJ, Reilly L, Treisman I, Cortés P, Coggiano MA, Veeravalli V, Rais R, Tanda G, Snyder SH (2021) Cocaine-induced locomotor stimulation involves autophagic degradation of the dopamine transporter. Mol Psychiatry 2021 262 26:370–382.

Harrod SB, Mactutus CF, Fitting S, Hasselrot U, Booze RM (2008) Intra-accumbal Tat1-72 alters acute and sensitized responses to cocaine. Pharmacol Biochem Behav 90:723.

Hong AR, Jang JG, Chung YC, Won SY, Jin BK (2022) Interleukin 13 on Microglia is Neurotoxic in Lipopolysaccharide-injected Striatum in vivo. Exp Neurobiol 31:42.

Huang YH, Lin Y, Mu P, Lee BR, Brown TE, Wayman G, Marie H, Liu W, Yan Z, Sorg BA, Schlüter OM, Zukin RS, Dong Y (2009) In vivo cocaine expearience generates silent synapses. Neuron 63:40–47.

Kalivas PW (2004) Glutamate systems in cocaine addiction. Curr Opin Pharmacol 4:23–29.

Kalivas PW (2009) The glutamate homeostasis hypothesis of addiction. Nat Rev Neurosci 10:561–572.

Kalivas PW, Duffy P (1990) Effect of acute and daily cocaine treatment on extracellular dopamine in the nucleus accumbens. Synapse 5:48–58.

Kau KS, Madayag A, Mantsch JR, Grier MD, Abdulhameed O, Baker DA (2008) Blunted cystine-glutamate antiporter function in the nucleus accumbens promotes cocaine-induced drug seeking. Neuroscience 155:530–537 Available at: https://www-clinicalkey-com.ezproxy2.library.drexel.edu/#!/content/playContent/1-s2.0-S0306452208008890?returnurl= https://linkinghub.elsevier.com%2Fretrieve%2Fpii%2FS0306452208008890%3Fshowall%3Dtrue&referrer= [Accessed March 2, 2020].

Kelschenbach J, He H, Kim BH, Borjabad A, Gu CJ, Chao W, Do M, Sharer LR, Zhang H, Arancio O, Potash MJ, Volsky DJ (2019) Efficient expression of HIV in immunocompetent mouse brain reveals a novel nonneurotoxic viral function in hippocampal synaptodendritic injury and memory impairment. MBio 10.

Kelschenbach JL, Saini M, Hadas E, Gu C, Chao W, Bentsman G, Hong JP, Hanke T, Sharer LR, Potash MJ, Volsky DJ (2012) Mice Chronically Infected with Chimeric HIV Resist Peripheral and Brain Superinfection: A Model of Protective Immunity to HIV. J Neuroimmune Pharmacol 7:380.

Kesby JP, Markou A, Semenova S (2016) The effects of HIV-1 regulatory TAT protein expression on brain reward function, response to psychostimulants and delay-dependent memory in mice. Neuropharmacology 109:205.

Kesby JP, Najera JA, Romoli B, Fang Y, Basova L, Birmingham A, Marcondes MCG, Dulcis D, Semenova S (2017) HIV-1 TAT protein enhances sensitization to methamphetamine by affecting dopaminergic function. Brain Behav Immun 65:210–221.

Kim R, Sepulveda-Orengo MT, Healey KL, Williams EA, Reissner KJ (2018) Regulation of glutamate transporter 1 (GLT-1) gene expression by cocaine self-administration and withdrawal. Neuropharmacology 128:1–10.

Knackstedt LA, Melendez RI, Kalivas PW (2010) Ceftriaxone Restores Glutamate Homeostasis and Prevents Relapse to Cocaine Seeking. Biol Psychiatry 67:81–84.

Koob GF, Volkow ND (2010) Neurocircuitry of addiction. Neuropsychopharmacology 35:217–238.

Koob GF, Volkow ND (2016) Neurobiology of addiction: a neurocircuitry analysis. The Lancet Psychiatry 3:760–773.

Koya E, Golden SA, Harvey BK, Guez-Barber DH, Berkow A, Simmons DE, Bossert JM, Nair SG, Uejima JL, Marin MT, Mitchell TB, Farquhar D, Ghosh SC, Mattson BJ, Hope BT (2009) Targeted disruption of cocaine-activated nucleus accumbens neurons prevents context-specific sensitization. Nat Neurosci 12:1069–1073.

Li Y, Acerbo MJ, Robinson TE (2004) The induction of behavioural sensitization is associated with cocaine-induced structural plasticity in the core (but not shell) of the nucleus accumbens. Eur J Neurosci 20:1647– 1654.

Liu X, Chang L, Vigorito M, Kass M, Li H, Chang SL (2009) Methamphetamine-induced behavioral sensitization is enhanced in the HIV-1 transgenic rat. J Neuroimmune Pharmacol 4:309–316.

Maragos WF, Young KL, Turchan JT, Guseva M, Pauly JR, Nath A, Cass WA (2002) Human immunodeficiency virus-1 Tat protein and methamphetamine interact synergistically to impair striatal dopaminergic function. J Neurochem 83:955–963.

McFarland K, Lapish CC, Kalivas PW (2003) Prefrontal glutamate release into the core of the nucleus accumbens mediates cocaine-induced reinstatement of drug-seeking behavior. J Neurosci 23:3531–3537.

McIntosh S, Sexton T, Pattison LP, Childers SR, Hemby SE (2015) Increased Sensitivity to Cocaine Self-Administration in HIV-1 Transgenic Rats is Associated with Changes in Striatal Dopamine Transporter Binding. J Neuroimmune Pharmacol 10:493–505.

McLaurin KA, Cook AK, Li H, League AF, Mactutus CF, Booze RM (2018) Synaptic connectivity in medium spiny neurons of the nucleus accumbens: A sex-dependent mechanism underlying apathy in the HIV-1 transgenic rat. Front Behav Neurosci 12.

Meade CS, Addicott M, Hobkirk AL, Towe SL, Chen NK, Sridharan S, Huettel SA (2018) Cocaine and HIV are independently associated with neural activation in response to gain and loss valuation during economic risky choice. Addict Biol 23:796.

Melendez RI, Roman C, Capo-Velez CM, Lasalde-Dominicci JA (2016) Decreased glial and synaptic glutamate uptake in the striatum of HIV-1 gp120 transgenic mice. J Neurovirol 22:358–365.

Mimiaga MJ, Reisner SL, Grasso C, Crane HM, Safren SA, Kitahata MM, Schumacher JE, Mathews WC, Mayer KH (2013) Substance use among HIV-infected patients engaged in primary care in the United States: Findings from the centers for AIDS Research Network of Integrated Clinical Systems Cohort. Am J Public Health 103:1457–1467.

Namba MD, Xie Q, Barker JM (2023) Advancing the preclinical study of comorbid neuroHIV and substance use disorders: Current perspectives and future directions. Brain Behav Immun 113:453–475.

Namba MD, Xie Q, Park K, Jackson JG, Barker JM (2024) EcoHIV Infection Modulates the Effects of Cocaine Exposure Pattern and Abstinence on Cocaine Seeking and Neuroimmune Protein Expression in Male Mice. bioRxiv Prepr Serv Biol.

Nickoloff-Bybel EA, Calderon TM, Gaskill PJ, Berman JW (2020) HIV Neuropathogenesis in the Presence of a Disrupted Dopamine System. J Neuroimmune Pharmacol 15:729–742.

Nooka S, Ghorpade A (2017) HIV-1-associated inflammation and antiretroviral therapy regulate astrocyte endoplasmic reticulum stress responses. Cell Death Discov 3.

O’Donovan B, Neugornet A, Neogi R, Xia M, Ortinski P (2021) Cocaine experience induces functional adaptations in astrocytes: implications for synaptic plasticity in the nucleus accumbens shell. Addict Biol 26:e13042.

Olson KE, Bade AN, Namminga KL, Potash MJ, Mosley RL, Poluektova LY, Volsky DJ, Gendelman HE (2018) Persistent EcoHIV infection induces nigral degeneration in 1-methyl-4-phenyl-1,2,3,6-tetrahydropyridine-intoxicated mice. J Neurovirol 24:398–410.

Pandey HS, Kapoor R, Bindu, Seth P (2022) Coronin 1A facilitates calcium mobilization and promotes astrocyte reactivity in HIV-1 neuropathogenesis. FASEB BioAdvances 4:254.

Pang S, Koyanagi Y, Miles S, Wiley C, Vinters H V., Chen ISY (1990) High levels of unintegrated HIV-1 DNA in brain tissue of AIDS dementia patients. Nature 343:85–89.

Paris J, Fenwick J, McLaughlin J (2015) Estrous Cycle and HIV-1 Tat Protein Influence Cocaine-Conditioned Place Preference and Induced Locomotion of Female Mice. Curr HIV Res 12:388–396.

Paris JJ, Carey AN, Shay CF, Gomes SM, He JJ, McLaughlin JP (2014) Effects of conditional central expression of HIV-1 tat protein to potentiate cocaine-mediated psychostimulation and reward among male mice. Neuropsychopharmacology 39:380–388.

Parsons LH, Justice JB (1993) Serotonin and dopamine sensitization in the nucleus accumbens, ventral tegmental area, and dorsal raphe nucleus following repeated cocaine administration. J Neurochem 61:1611–1619.

Perrine SA, Ghoddoussi F, Desai K, Kohler RJ, Eapen AT, Lisieski MJ, Angoa-Perez M, Kuhn DM, Bosse KE, Conti AC, Bissig D, Berkowitz BA (2015) Cocaine-induced locomotor sensitization in rats correlates with nucleus accumbens activity on manganese-enhanced MRI. NMR Biomed 28:1480–1488.

Pittaluga A (2017) CCL5-glutamate cross-talk in astrocyte-neuron communication in multiple sclerosis. Front Immunol 8:285787.

Potash MJ, Chao W, Bentsman G, Paris N, Saini M, Nitkiewicz J, Belem P, Sharer L, Brooks AI, Volsky DJ (2005) A mouse model for study of systemic HIV-1 infection, antiviral immune responses, and neuroinvasiveness. Proc Natl Acad Sci U S A 102:3760–3765.

Reissner KJ, Gipson CD, Tran PK, Knackstedt LA, Scofield MD, Kalivas PW (2015) Glutamate transporter GLT-1 mediates N-acetylcysteine inhibition of cocaine reinstatement. Addict Biol 20:316–323.

Riday TT, Kosofsky BE, Malanga CJ (2012) The Rewarding and Locomotor-Sensitizing Effects of Repeated Cocaine Administration are Distinct and Separable in Mice. Neuropharmacology 62:1858–1866.

Satarker S, Bojja SL, Gurram PC, Mudgal J, Arora D, Nampoothiri M (2022) Astrocytic Glutamatergic Transmission and Its Implications in Neurodegenerative Disorders. Cells 11.

Scofield MD, Boger HA, Smith RJ, Li H, Haydon PG, Kalivas PW (2015) Gq-DREADD selectively initiates glial glutamate release and inhibits cue-induced cocaine seeking. Biol Psychiatry 78:441–451.

Scofield MD, Li H, Siemsen BM, Healey KL, Tran PK, Woronoff N, Boger HA, Kalivas PW, Reissner KJ (2016) Cocaine Self-Administration and Extinction Leads to Reduced Glial Fibrillary Acidic Protein Expression and Morphometric Features of Astrocytes in the Nucleus Accumbens Core. Biol Psychiatry 80:207–215.

Siemsen BM, Reichel CM, Leong KC, Garcia-Keller C, Gipson CD, Spencer S, McFaddin JA, Hooker KN, Kalivas PW, Scofield MD (2019) Effects of Methamphetamine Self-Administration and Extinction on Astrocyte Structure and Function in the Nucleus Accumbens Core. Neuroscience 406:528–541.

Sofroniew M V., Vinters H V. (2010) Astrocytes: Biology and pathology. Acta Neuropathol 119:7–35.

Steketee JD, Kalivas PW (2011) Drug wanting: Behavioral sensitization and relapse to drug-seeking behavior. Pharmacol Rev 63:348–365.

Sun W, Cornwell A, Li J, Peng S, Joana Osorio M, Aalling N, Wang S, Benraiss A, Lou N, Goldman SA, Nedergaard M (2017) SOX9 Is an Astrocyte-Specific Nuclear Marker in the Adult Brain Outside the Neurogenic Regions. J Neurosci 37:4493.

Teo I, Veryard C, Barnes H, An SF, Jones M, Lantos PL, Luthert P, Shaunak S (1997) Circular forms of unintegrated human immunodeficiency virus type 1 DNA and high levels of viral protein expression: association with dementia and multinucleated giant cells in the brains of patients with AIDS. J Virol 71:2928.

Thomas MJ, Beurrier C, Bonci A, Malenka RC (2001) Long-term depression in the nucleus accumbens: A neural correlate of behavioral sensitization to cocaine. Nat Neurosci 4:1217–1223.

Van Zessen R, Li Y, Marion-Poll L, Hulo N, Flakowski J, Lüscher C (2021) Dynamic dichotomy of Accumbal population activity underlies cocaine sensitization. Elife 10.

Wallet C, De Rovere M, Van Assche J, Daouad F, De Wit S, Gautier V, Mallon PWG, Marcello A, Van Lint C, Rohr O, Schwartz C (2019) Microglial Cells: The Main HIV-1 Reservoir in the Brain. Front Cell Infect Microbiol 9:362.

Wang J, Holt LM, Huang HH, Sesack SR, Nestler EJ, Dong Y (2021a) Astrocytes in cocaine addiction and beyond. Mol Psychiatry:1–17.

Wang J, Li KL, Shukla A, Beroun A, Ishikawa M, Huang X, Wang Y, Wang YQ, Yang Y, Bastola ND, Huang HH, Kramer LE, Chao T, Huang YH, Sesack SR, Nestler EJ, Schlüter OM, Dong Y (2021b) Cocaine Triggers Astrocyte-Mediated Synaptogenesis. Biol Psychiatry 89:386–397.

Wang Z, Pekarskaya O, Bencheikh M, Chao W, Gelbard HA, Ghorpade A, Rothstein JD, Volsky DJ (2003) Reduced expression of glutamate transporter EAAT2 and impaired glutamate transport in human primary astrocytes exposed to HIV-1 or gp120. Virology 312:60–73.

Xie Q, Namba MD, Buck LA, Park K, Jackson JG, Barker JM (2024) Effects of Antiretroviral Treatment on Central and Peripheral Immune Response in Mice with EcoHIV Infection. Cells 13:882.

